# Expanding the toolkit for genetic manipulation and discovery in *Candida* species using a CRISPR ribonucleoprotein-based approach

**DOI:** 10.1101/2023.06.16.545382

**Authors:** Justin B. Gregor, Victor A. Gutierrez-Schultz, Smriti Hoda, Kortany M. Baker, Debasmita Saha, Madeline G. Burghaze, Scott D. Briggs

**Affiliations:** Department of Biochemistry and; Purdue University Institute for Cancer Research

**Keywords:** CRISPR, *Candida auris*, ribonucleoprotein particle (RNP), *Candida glabrata*, *Candida albicans*, genome editing, drug resistance cassette, homology directed repair, fungal pathogens, *KanMX* and *BleMX*, ergosterol pathway, Set1 histone H3K4 methyltransferase

## Abstract

The World Health Organization recently published the first list of priority fungal pathogens highlighting multiple *Candida* species including *C. glabrata*, *C. albicans*, and *C. auris*. The use of CRISPR-Cas9 and auxotrophic *C. glabrata* and *C. albicans* strains have been instrumental in the study of these fungal pathogens. Dominant drug resistance cassettes are also critical for genetic manipulation and eliminate the concern of altered virulence when using auxotrophic strains. However, genetic manipulation has been mainly limited to the use of two drug resistance cassettes, *NatMX* and *HphMX*. Using an *in vitro* assembled CRISPR-Cas9 ribonucleoprotein (RNP)-based system and 130-150 bp homology regions for directed repair, we expand the drug resistance cassettes for *Candida* to include *KanMX* and *BleMX*, commonly used in *S. cerevisiae*. As a proof of principle, we demonstrated efficient deletion of *ERG* genes using *KanMX* and *BleMX*. We also showed the utility of the CRISPR-Cas9 RNP system for generating double deletions of genes in the ergosterol pathway and endogenous epitope tagging of *ERG* genes using an existing *KanMX* cassette. This indicates that CRISPR-Cas9 RNP can be used to repurpose the *S. cerevisiae* toolkit. Furthermore, we demonstrated that this method is effective at deleting *ERG3* in *C. auris* using a codon optimized *BleMX* cassette and effective at deleting the epigenetic factor, *SET1*, in *C. albicans* using a recyclable *SAT1.* Using this expanded toolkit, we discovered new insights into fungal biology and drug resistance.

**IMPORTANCE:** The increasing problem of drug resistance and emerging pathogens is an urgent global health problem that necessitates the development and expansion of tools for studying fungal drug resistance and pathogenesis. We have demonstrated the effectiveness of an expression-free CRISPR-Cas9 RNP-based approach employing 130-150 bp homology regions for directed repair. Our approach is robust and efficient for making gene deletions in *C. glabrata*, *C. auris* and *C. albicans* as well as epitope tagging in *C. glabrata*. Furthermore, we demonstrated that *KanMX* and *BleMX* drug resistance cassettes can be repurposed in *C. glabrata* and *BleMX* in *C. auris*. Overall, we have expanded the toolkit for genetic manipulation and discovery in fungal pathogens.

## INTRODUCTION

Fungal infections pose a significant public health concern, with over a billion superficial infections and 1.5 million deaths occurring annually worldwide (1, 2). *Candida* species are responsible for roughly 40-70% of invasive fungal infections (1–3), and several species are classified as “high priority fungal pathogens” by the World Health Organization (WHO) for study, including *C. glabrata*, *C. albicans*, and *C. auris*. Infections can range from superficial to life-threatening, with invasive candidiasis leading to a mortality rate of 20-60% (4, 5). Currently, there are three major antifungals clinically used for treatment of fungal infections; azoles, echinocandins, and polyenes (6–8). However, antifungal drug resistance has become a significant concern, highlighted by the increase in clinically acquired drug resistance in *C. albicans* and *C. glabrata* and the recent emergence of a multi-drug resistant pathogen, *C. auris* (8, 9). With increased drug resistance and emerging pathogens, there is an urgent need for the development and expansion of new and existing tools for studying drug resistance and pathogenesis in *Candida*, especially in non-*albicans Candida* (NAC) species.

To address this need, several groups use various <u>C</u>lustered <u>R</u>egular Interspaced <u>S</u>hort <u>P</u>alindromic <u>R</u>epeats (CRISPR) based methods for genetic manipulations. CRISPR-Cas9 is a tool that utilizes the Cas9 endonuclease to direct double stranded breaks (DSBs) at the desired locus by binding to a gene specific guide RNA followed by a protospacer-adjacent motif (PAM) equence. A Cas9-mediated DSB will activate either the nonhomologous end-joining (NHEJ) for error-prone repair resulting in insertions or deletions or a precise homology directed repair (HDR) using a donor template. Using both approaches can greatly enhance the efficiency to generate genetic mutations, gene replacements, or epitope tags.

There are two common strategies for utilizing CRISPR genome editing in *C. albicans* or *C. glabrata*. One approach involves the expression of the Cas9 enzyme and sgRNA from separate plasmids while the other approach uses one plasmid for expressing Cas9 and sgRNA (10, 11). These plasmid-based approaches can be either episomally expressed or integrated in the genome (12, 13). Another approach is using an expression-free CRISPR-Cas9 ribonucleoprotein (RNP) method (10, 14–18). The CRISPR-Cas9 RNP approach has been used with a HDR template containing the *NAT1, SAT, and HygB* resistances cassette for generating gene deletions in *C. glabrata*, *C. auris*, *C. lusitaniae, and C. albicans* (14–18). A major advantage of this system is that the CRISPR-based RNP system does not require plasmid engineering or species-specific promoter expression in cells (10, 11, 18). Instead, recombinant Cas9 protein, crRNA and tracrRNA are assembled as a ribonucleoprotein complex *in vitro* and electroporated into competent cells which reduces the steps needed to genetically manipulate prototrophic strains or clinical isolates.

In this study, we used an expression-free CRISPR-Cas9 RNP-based approach using homology regions of 130-150 bp for making efficient gene deletions in *C. glabrata* and *C. auris*. Our CRISPR-Cas9 RNP approach showed improved efficiency over a non-CRISPR based method using *ADE2* as our reporter for gene disruption. Furthermore, all drug resistance cassettes used for gene disruptions and epitope tagging were PCR amplified using homology regions of 130-150 bp, indicating large flanking sequences are not required with the CRISPR- Cas9 RNP-based approach. Our approach also permits making double deletions and epitope tagging which are difficult to make without CRISPR. More importantly, we demonstrated the utilization of drug resistance cassettes *KanMX and BleMX* in *C. glabrata* for generating gene deletions and *KanMX* for generating epitope tags. These two drug resistance cassettes have not been widely used for *C. glabrata* but are extensively used for *S. cerevisiae*. Finally, we demonstrated that the CRISPR-Cas9 RNP approach can also be utilized for making gene deletions in *C. auris* using codon optimized *BleMX*. Overall, using the CRISPR-Cas9 RNP approach allowed us to expand the fungal pathogen toolkit by demonstrating that *KanMX* containing plasmids used for *S. cerevisiae* can be repurposed for *C. glabrata* and that *BleMX* can be used for *C. auris* and *C. glabrata*. Furthermore, we showed the utility of these tools by providing phenotypic characterization of factors that alter ergosterol biosynthesis and fluconazole drug susceptibility.

## RESULTS

### The CRISPR-Cas9 RNP system for efficient gene replacement in C. glabrata using 130- 150 bp homology regions

CRISPR-mediated or non-CRISPR-based methods generally rely on large flanking homologous regions ranging from 500 bp to 1000 bp for efficient gene replacements in *Candida* species (18, 19). Often steps to generate long flanking regions are time consuming and tedious using either cloning or multi-step fusion PCR approaches. The initial CRISPR-Cas9 RNP system developed for *Candida* species including *C. glabrata* utilized long homology regions ranging from 500 to 1000 bp (18). However, using CRISPR-Cas9 plasmid-based system and auxotrophic cassettes, it has been reported for *C. glabrata* that flanking homology regions ranging from 20-200 bp can be used for gene insertions resulting in gene disruption (12). To determine if short homology regions flanking drug resistance cassettes were efficient in making gene deletions in *C. glabrata* using a CRISPR-Cas9 RNP method, we PCR amplified drug resistance cassettes using long oligonucleotides (IDT Ultramers) that range from ∼130-150 bp of homology to the *ADE2* gene. The *ADE2* gene was selected due to its red pigment phenotype when *ADE2* gene is disrupted which allows for quick determination of gene replacement efficiency and has been commonly used to determine CRISPR efficiency (13, 20, 21). Using the pAG25 *NatMX* and pAG32 *HphMX* plasmids (Fig. 1A, 1B) (22), we deleted the entire *ADE2* open reading frame and counted the proportion of white and red colonies (Fig. 1C). With the addition of a CRISPR-Cas9 RNP containing two gRNAs and 130-150 bp of flanking homology, we observed a five-fold increase in the proportion of red colonies compared to the cassette alone (Fig. 1D). With an efficiency of 62% red colonies, we determined that long homology regions are not required for efficient gene replacement in *C. glabrata* using *NatMX*. Similarly, we observed a five-fold increase in the proportion of red colonies using Hygromycin B (*HphMX*), with an efficiency of 55% replacement using our CRISPR-Cas9 RNP method (Fig. 1E).

**FIG 1.**
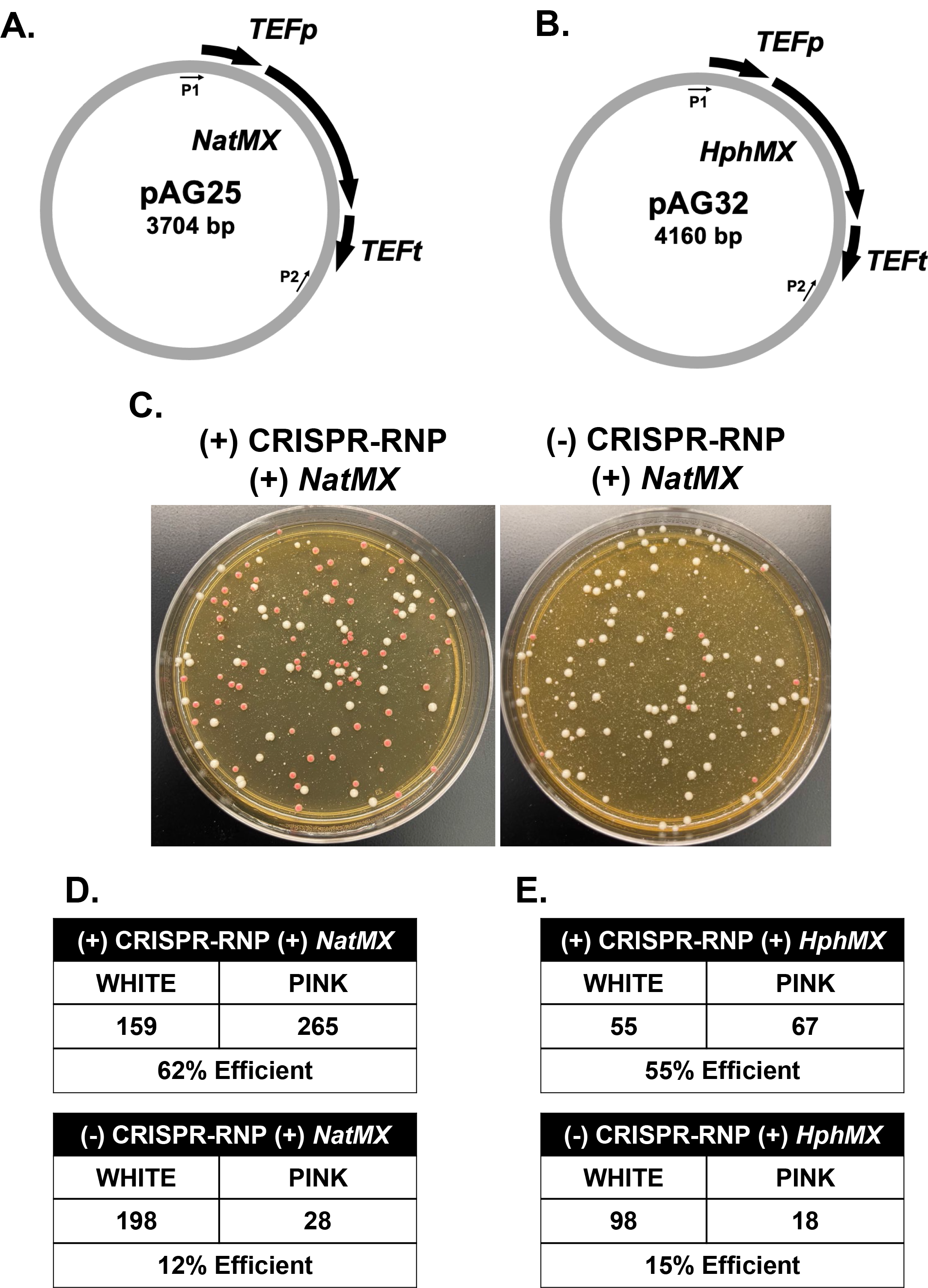
The CRISPR-Cas9 RNP system using 130-150 bp homology regions efficiently generates *ADE2* deletions in *C. glabrata* using *NatMX* and *HphMX*. **(A)** Schematic of pAG25 *NatMX* plasmid. P1 and P2 indicate location of amplification sequences. **(B)** Schematic of pAG32 *HphMX* plasmid. P1 and P2 indicate location of amplification sequences. **(C)** Representative transformation plate for *ADE2* deletion using *NatMX* with and without addition of CRISPR-RNP. **(D)** Total number of positive transformants using *NatMX* with and without addition of CRISPR-RNP. Numbers represent the summation across three separate transformations. **(E)** Total number of positive transformants using *HphMX* with and without addition of CRISPR-RNP. Numbers represent the summation across three separate transformations.

Altogether, these data suggest that our modified CRISPR-RNP method using 130-150 bps of homology can efficiently generate single gene deletions in *C. glabrata*.

### Using CRISPR-Cas9 RNP system to generate sequential gene replacements utilizing NatMX and HphMX *resistance cassettes*

After determining this system efficiently generates single gene deletions, we then tested whether the CRISPR-Cas9 RNP method was sufficient for making sequential gene disruptions for generating double deletion strains. While the pAG25 and pAG32 plasmids are effective for use in single deletions, it is often difficult to generate double gene deletions with these replacement cassettes using standard transformation methods. We suspect that making double deletions using drug resistance cassettes are difficult because they often share homology with the *TEF1* promoter and *TEF1* terminator (Fig. 1A, 1B). To overcome these issues, we tested if our CRISPR-Cas9 RNP method was sufficient for generating double gene deletions using *HphMX* and *NatMX* resistance cassettes.

For proof of principle, we probed the ergosterol biosynthesis pathway, a critical pathway for azole antifungal drugs. Azole drugs inhibit Erg11, lanosterol 14-alpha-demethylase, to block ergosterol biosynthesis and leads to accumulation of an Erg3-dependent toxic sterol 14α- methyl-3,6-diol and growth arrest (23–25). Thus, *ERG* gene deletions or mutations in this pathway can alter azole susceptibility and growth. For example, *ERG3* is known to have an azole resistant phenotype when deleted or mutated in *S. cerevisiae* or *C. albicans* which is a consequence of not producing the toxic sterol 14α-methyl-3,6-diol (24, 26, 27). However, there has been conflicting results in *C. glabrata* where *ERG3* deletions do not confer resistance to fluconazole while microevolved *ERG3* mutations and clinical isolates have shown resistance to fluconazole (25, 28–31). To address this issue, we used our CRISPR-Cas9 RNP method to make *erg3Δ* strains using the pAG25-*NatMX* and pAG32-*HphMX* as a template or *erg5Δ* strain using *NatMX* (Fig. 1A, B). We performed five-fold serial dilution spot assays on these strains to confirm and compare their phenotypes with and without 64 μg/mL fluconazole in SC media (Fig. 2A, B). Both *erg3Δ* strains demonstrate a slow growth phenotype, but also a clear increased resistance to fluconazole (Fig. 2A). However, the *erg5Δ* strain did not have an observable growth defect without fluconazole and little to no growth on fluconazole containing plates (Fig. 2B).

**FIG 2.**
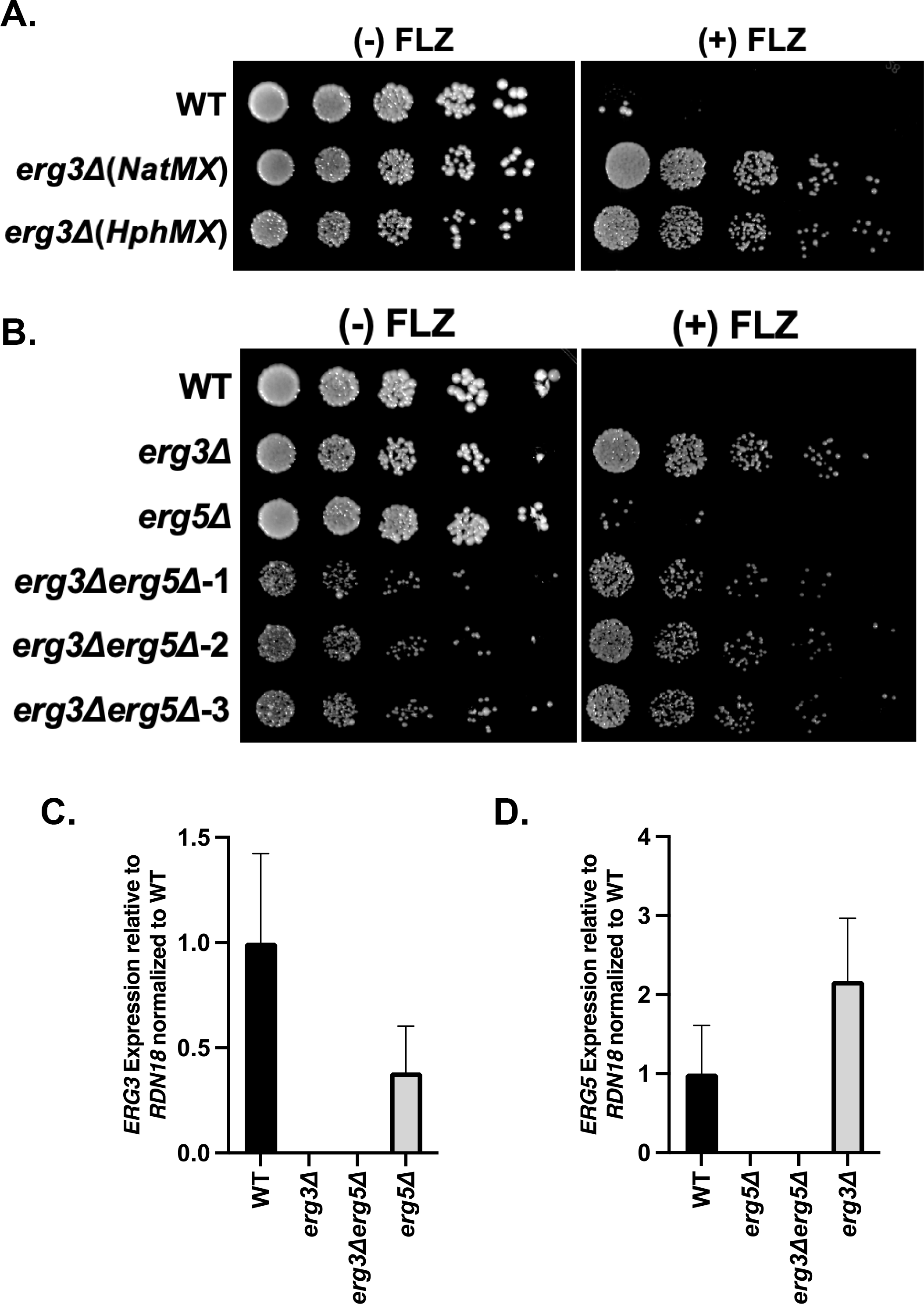
The CRISPR-Cas9 RNP system generates single and double gene deletions utilizing *NatMX* and *HphMX* in *C. glabrata*. **(A and B)** Five-fold serial dilution spot assays with and without 64 μg/mL fluconazole (FLZ). Indicated single deletion strains were generated using the CRISPR-Cas9 RNP method. Double deletion strains were generated using CRISPR-Cas9 RNP method sequentially and three independent clones are shown. Images were captured at 48 hours. **(C and D)** Expression of the indicated genes were determined by qRT-PCR analysis of mid-log phase cells. Data was normalized to *RDN18* mRNA levels and are the average of three biological replicates with three technical replicates each. Error bars represent the standard deviation.

Next, we used our CRISPR-Cas9 RNP method to generate *erg3Δerg5Δ* double deletion strains, by deleting *ERG5* with *HphMX* in the previously constructed *erg3Δ* (*NatM*X) strain. After confirming positive transformants via colony PCR, five-fold serial dilution spot assays with and without 64 μg/mL fluconazole were performed. Interestingly, all *erg3Δerg5Δ* strains display a synthetic growth defect, more than what was observed in the single *erg3Δ* and *erg5Δ* strains (Fig. 2B). Despite this significant growth defect under untreated conditions, *erg3Δerg5Δ* strains were still able to grow on fluconazole containing plates similar to an *erg3Δ* strain (Fig. 2B). To further validate these strains, we grew cells in SC media to mid-log phase and collected cells for qRT-PCR expression on both *ERG3* and *ERG5*. In each strain lacking *ERG3*, we detected no *ERG3* transcript, confirming that *ERG3* was deleted (Fig. 2C). Additionally, in each strain lacking *ERG5*, we detected no *ERG5* expression (Fig, 2D), confirming *ERG5* was deleted.

Interestingly, we do see decreased expression of *ERG3* in the *erg5Δ* strain and increased expression of *ERG5* in the *erg3Δ* strain which is consistent with what is observed in *S. cerevisiae* (32, 33). Altogether, these results suggest that our CRISPR-Cas9 RNP method permits engineering of single and double deletions in *C. glabrata*. Moreover, we clearly established that *erg3Δ* strains are resistant to fluconazole and identify a genetic interaction between *ERG3 and ERG5*.

### The CRISPR-Cas9 RNP system efficiently generates gene deletions utilizing the BleMX drug resistance cassette in C. glabrata

Since our CRISPR-Cas9 RNP system is efficient at generating single and double deletions in *C. glabrata* using *NatMX* and *HphMX*, we then tested whether this system was effective for using other drug resistance cassettes typically not used in *C. glabrata*. We first tested *BleMX*, which confers resistance to Zeocin, as the use of *BleMX* has been reported once in *C. glabrata* using a non-CRISPR transformation method, albeit at extremely low efficiency (<1%) (34). To first determine whether the CRISPR-Cas9 RNP system effectively generates gene deletions using *BleMX*, we deleted the entire open reading frame of *ADE2* using the pCY3090-07 plasmid as a template (Fig. 3A) (35). When comparing the proportion of red colonies with and without the addition of CRISPR-Cas9, a five to six-fold increase in efficiency was observed (Fig. 3B). Next, using the pCY3090-07 plasmid, we deleted *ERG3* using our CRISPR-Cas9 RNP method and subsequently performed a five-fold serial dilution spot assay with and without 64 μg/mL fluconazole to compare phenotypes of these strains with previously constructed *erg3Δ* strains. Similar to the previously constructed *erg3Δ* strains, we again observed an azole resistant phenotype. Thus, our results using 4 different drug resistance cassettes clearly indicate that *erg3Δ* strains are resistant to fluconazole under the indicated conditions (Fig. 3C and 4C). These data show that *BleMX* is an effective dominant drug resistance cassette in *C. glabrata* when using the CRISPR-Cas9 RNP system.

**FIG 3.**
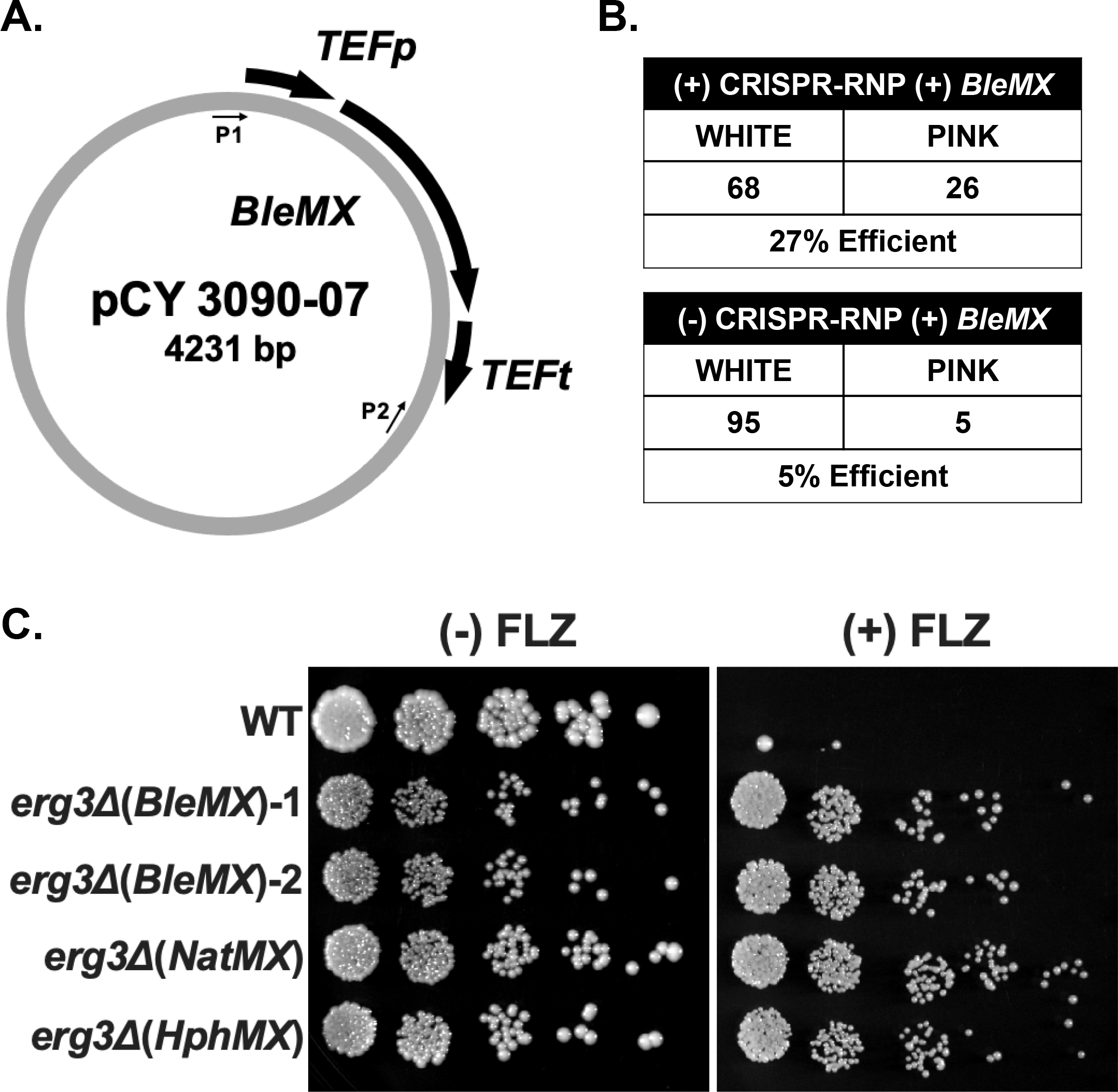
The CRISPR-Cas9 RNP system efficiently generates gene deletions utilizing *BleMX* in *C. glabrata*. **(A)** Schematic of pCY3090-07 plasmid. P1 and P2 indicate location of amplification primer sequences. **(B)** Total number of positive transformants using *BleMX* with and without addition of CRISPR-Cas9 RNP. Numbers are the summation across three separate transformations. **(C)** Five-fold serial dilution spot assays of indicated strains with and without 64 μg/mL fluconazole (FLZ). Two independent clones are shown for *erg3Δ* (*BleMX*). Images were captured at 48 hours.

**FIG 4.**
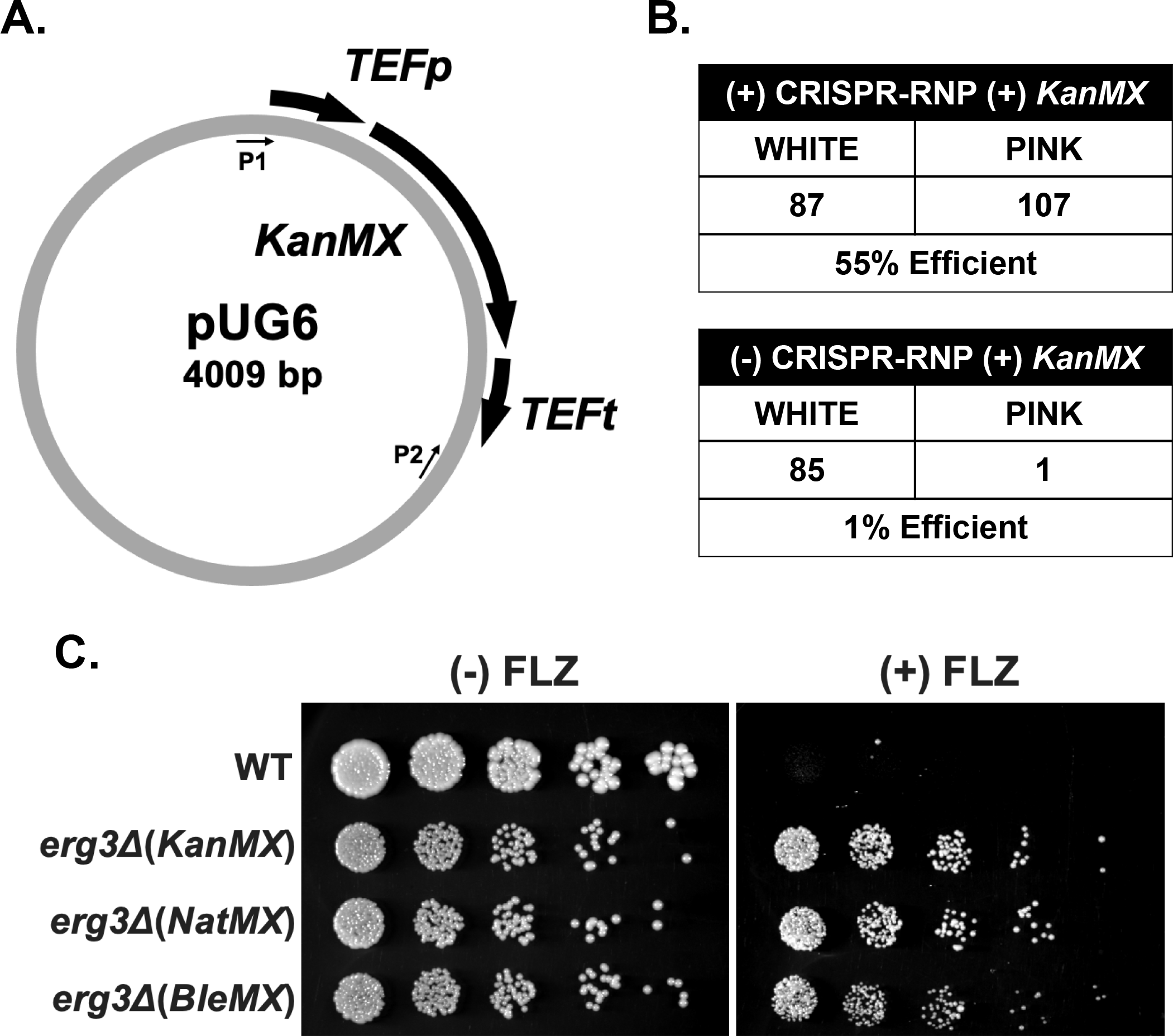
The CRISPR-RNP system efficiently generates gene deletions utilizing *KanMX* for *C. glabrata*. **(A)** Schematic of pUG6 plasmid. P1 and P2 indicate location of amplification primer sequences. **(B)** Total number of positive transformants using *KanMX* with and without addition of CRISPR-RNP. Numbers are the summation across three separate transformations. **(C)** Five- fold serial dilution spot assays of indicated strains with and without 64 μg/mL fluconazole (FLZ). Images were captured at 48 hours.

### The CRISPR-Cas9 RNP system efficiently generates gene deletions utilizing the KanMX drug resistance cassette in C. glabrata

After determining that our CRISPR-Cas9 RNP was effective for single and double gene deletions using *NatMX*, *HphMX* and *BleMX*, we tested whether this system permitted the use of *KanMX* as a drug resistance cassette in *C. glabrata*. Although *KanMX* is routinely used in *S. cerevisiae*, *KanMX* has not been successfully utilized for genetic manipulations in *C. glabrata*. The use of *KanMX* drug resistance cassettes would allow the repurposing of many *S. cerevisiae* tagging and deletion *KanMX* cassettes for *C. glabrata*. In our studies, we have successfully generated several *C. glabrata* deletion strains using *HphMX* or *NatMX* using ∼130-150bp homology with chemical transformation and electroporation (36, 37). However, any attempts to use *KanMX* as a drug selection cassette using these two methods were not successful (data not shown and Fig. 4B). To first determine whether the CRISPR-Cas9 RNP system is sufficient for repurposing *KanMX* for use in *C. glabrata*, *ADE2* was deleted using the *KanMX* drug resistance cassette amplified from the pUG6 plasmid (Fig. 4A) (38). We observed 55% red colonies suggesting that the CRISPR-Cas9 RNP method is efficient in generating gene deletions using *KanMX* and 800 µg/ml G418 (Fig. 4B). In contrast, when using electroporation without the aid of CRISPR-Cas9, only one red colony out of 86 was observed indicating an efficiency of 1.1%. We were also able to successfully delete *ERG3* with our CRISPR-Cas9 RNP method using *KanMX*. A five-fold serial dilution spot assay with and without 64 μg/mL fluconazole were performed to compare the phenotype to previously constructed *erg3Δ* strains. Importantly, we observe an azole resistant phenotype similar to the other constructed *erg3Δ* strains (Fig. 4C). These data demonstrate that *KanMX* is an effective drug resistance cassette for use in *C. glabrata* when using the CRISPR-Cas9 RNP approach.

### *The* CRISPR-RNP system generates endogenous epitope tagged proteins using KanMX in C. glabrata

Because our data indicate that *KanMX* is a suitable drug resistance cassette for gene deletions in *C. glabrata*, we next determined if using the CRISPR-Cas9 RNP method would also permit endogenous epitope tagging using the C-terminal 3xHA-*KanMX* plasmid (pFA6a) commonly used for *S. cerevisiae* (Fig. 5A) (39). *ERG3* and *ERG11* were used to demonstrate that the CRISPR-Cas9 RNP method would allow for endogenous C-terminal tagging using *KanMX*. After confirming the presence of the insert via colony PCR, strains were grown with and without 64 μg/mL fluconazole in SC media and collected at mid-log phase for immunoblotting using anti-HA (12CA5). Histone H3 was used as a loading control. Our data indicate that Erg3 protein is expressed under untreated conditions and induced under fluconazole treatment (Fig. 5B), which is consistent with transcript analysis from our previous study (40). Erg11 protein is also expressed under untreated conditions and induced under fluconazole treatment (Fig. 5C) similar to what is observed for Erg3 (Fig. 5B) and consistent with previous transcript and protein analysis (40, 41). To confirm that the epitope tag does not alter azole susceptibility, we performed five-fold serial dilution spot assays with and without 64 μg/mL fluconazole, using an *erg3Δ* strain as a control. All epitope tagged Erg3-3xHA strains grow similar to WT under both untreated and fluconazole treatment (Fig. 5D). We observe the same effect for Erg11-3xHA strains with and without fluconazole treatment (Fig. 5E). Altogether, these data suggest that the CRISPR-Cas9 RNP system effectively generates endogenous epitope tagged proteins using *KanMX* and C-terminally tagging Erg11 and Erg3 does not appear to alter azole susceptibility.

**FIG 5.**
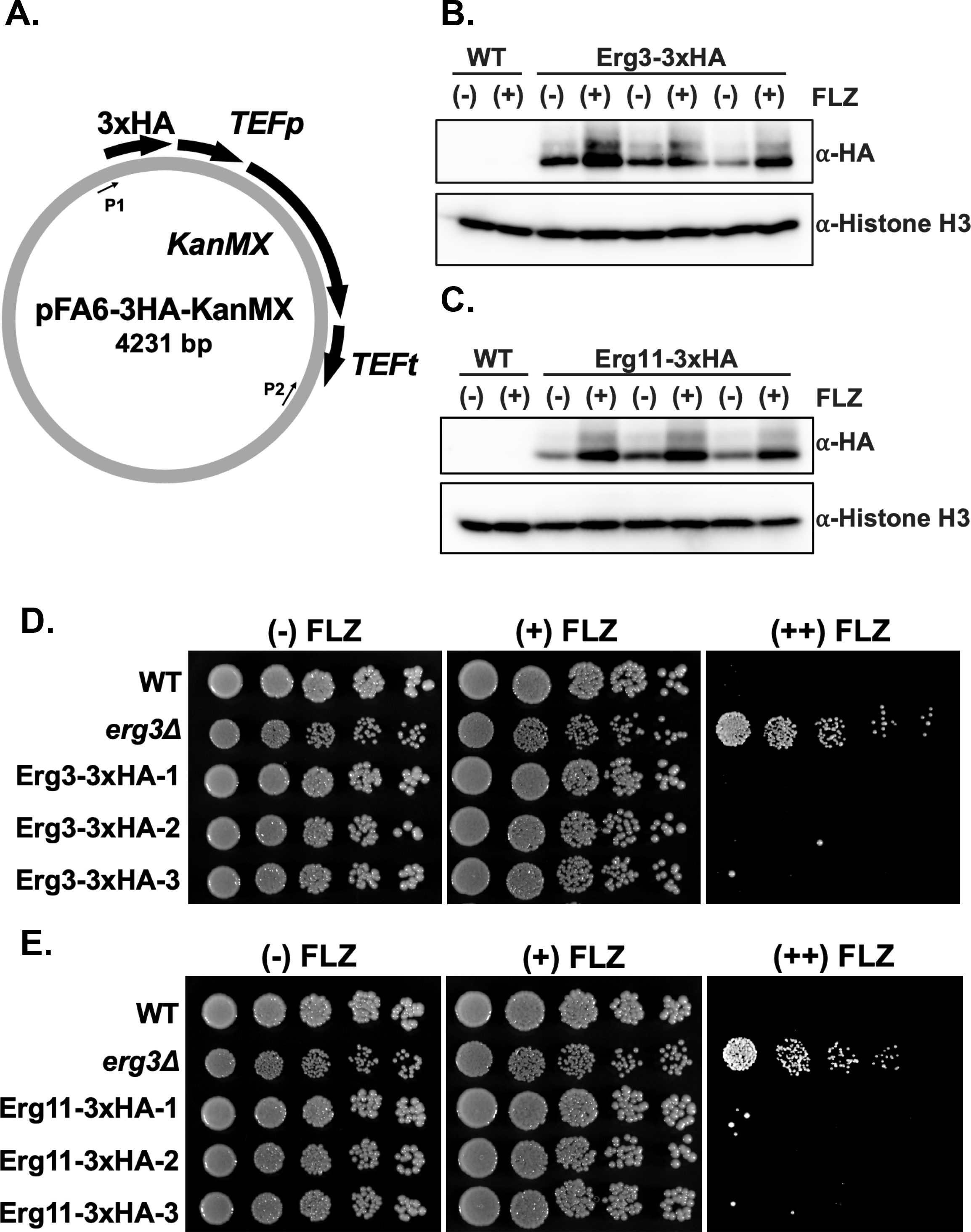
The CRISPR-Cas9 RNP system generates endogenous epitope tagged proteins using *KanMX* in *C. glabrata*. **(A)** Schematic of pFA6-3HA-KanMX plasmid. P1 and P2 indicate location of amplification primer sequences. **(B and C)** Indicated strains were either untreated (-) or treated (+) with 64 μg/mL of fluconazole (FLZ) for three hours. Whole cell extracts were isolated and immunoblotted against HA antibody for detection of Erg3 or Erg11. Histone H3 was used as a loading control. Three independent clones were represented for Erg3-3xHA and Erg11-3xHA. **(D and E)** Five-fold serial dilution spot assays of indicated strains with 0, 16, and 64 μg/mL fluconazole (FLZ), respectively. Three independent clones were represented for Erg3-3xHA and Erg11-3xHA. Images were captured at 48 hours.

### The CRISPR-Cas9 RNP system allows the use of BleMX as a drug resistance cassette for ***C. auris.***

It has been previously demonstrated that a CRISPR-Cas9 RNP system can be utilized in *C. auris* using *SAT1* as a drug resistance cassette (14–16, 18). With this, we tested whether the CRISPR-Cas9 RNP system allowed for the utilization of *BleMX* as a drug resistance cassette in *C. auris*. We first codon-optimized *BleMX* for use in CTG-clade species and named the plasmid pCdOpt-BMX (Fig. 6A). Using this codon-optimized *BleMX* plasmid as a template, we deleted *ERG3* in *C. auris AR0387* using the CRISPR-Cas9 RNP method. After confirming the presence of *BleMX* via colony PCR, we performed five-fold serial dilution spot assays with and without 64 μg/mL fluconazole on each strain. Similar to the *C. glabrata erg3Δ* strains, we observed a similar azole resistant phenotype across all clones (Fig. 6B). These data determine for the first time the effective use of our codon optimized *BleMX* in *C. auris* when using the CRISPR-RNP approach.

**FIG 6.**
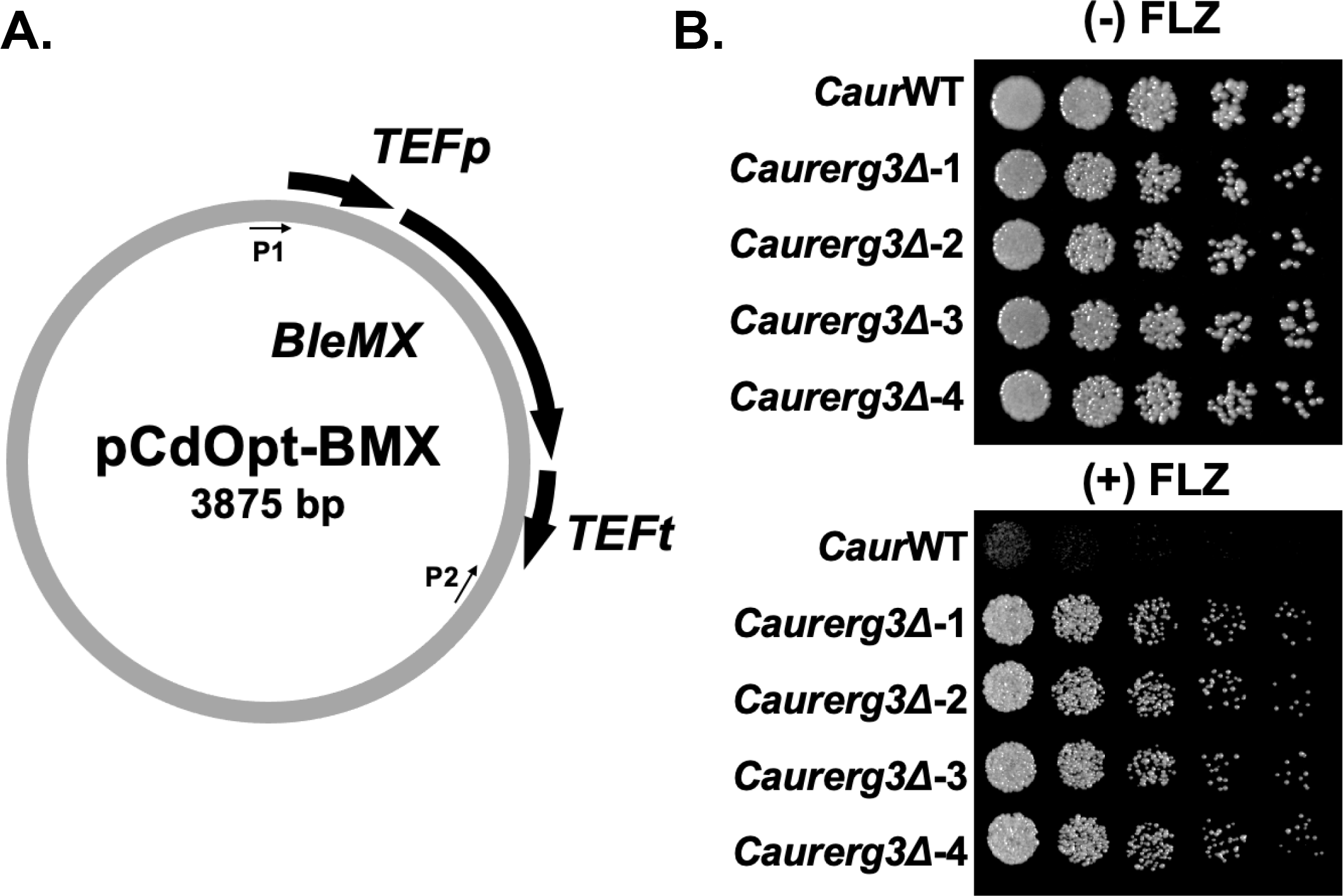
The CRISPR-Cas9 RNP system generates gene deletions using a codon optimized *BleMX* in *C. auris*. **(A)** Schematic of pCdOpt-BMX plasmid. P1 and P2 indicate location of amplification primer sequences. **(B)** Five-fold serial dilution spot assays of indicated *C. auris* strains with and without 64 μg/mL fluconazole (FLZ). Four independent clones were represented for *Caurerg3Δ* strain (*BleMX*). Images were captured at 48 hours.

### Using CRISPR-Cas9 RNP system to generate heterozygous and homozygous deletions in *C. albicans*

*SET1*, a known histone methyltransferase, when deleted in *C. glabrata* or *S. cerevisiae* alters azole susceptibility and *ERG* gene expression including *ERG3* (37, 42). We sought to determine whether loss of *SET1* in *C. albicans* exhibits a similar phenotype. To test this, we generated both heterozygous and homozygous *SET1* deletion mutants in a sequential manner where the entire open reading frame was deleted with the *SAT1* selection marker to generate the heterozygous deletion and then subsequently recycled using FLP recombinase to make the homozygous deletion in the *C. albicans* SC5314 strain (Fig. 7A) (43). Because Set1 is a histone H3K4 methyltransferase, we wanted to confirm the loss of methylation in the *set1Δ/Δ* strains using immunoblot analysis using H3K4me1, H3K4me2, and H3K4me3 specific antibodies (Fig. 7B). Since Set1 is the sole histone H3K4 methyltransferase in most yeast species, we observed a complete loss of H3K4 methylation in the *set1Δ/Δ* strain which is consistent with previous reports in a CAI4 strain (44, 45). We also determined that the *SET1/set1Δ* strain exhibited no change in H3K4 methylation status indicating loss of one allele didn’t impact global histone methylation (Fig. 7B). To assess azole susceptibility in *C. albicans*, we performed five-fold serial dilution spot assays with and without 0.5 µg/mL fluconazole. Interestingly, we did not observe altered susceptibility to azoles in the *set1Δ/Δ* or *SET1/set1Δ* strains (Fig. 7C). This is in clear contrast to what is observed when *SET1* is deleted in *C. glabrata* and *S. cerevisiae* indicating a species-specific difference and utilization of *SET1* (36, 37).

**FIG 7.**
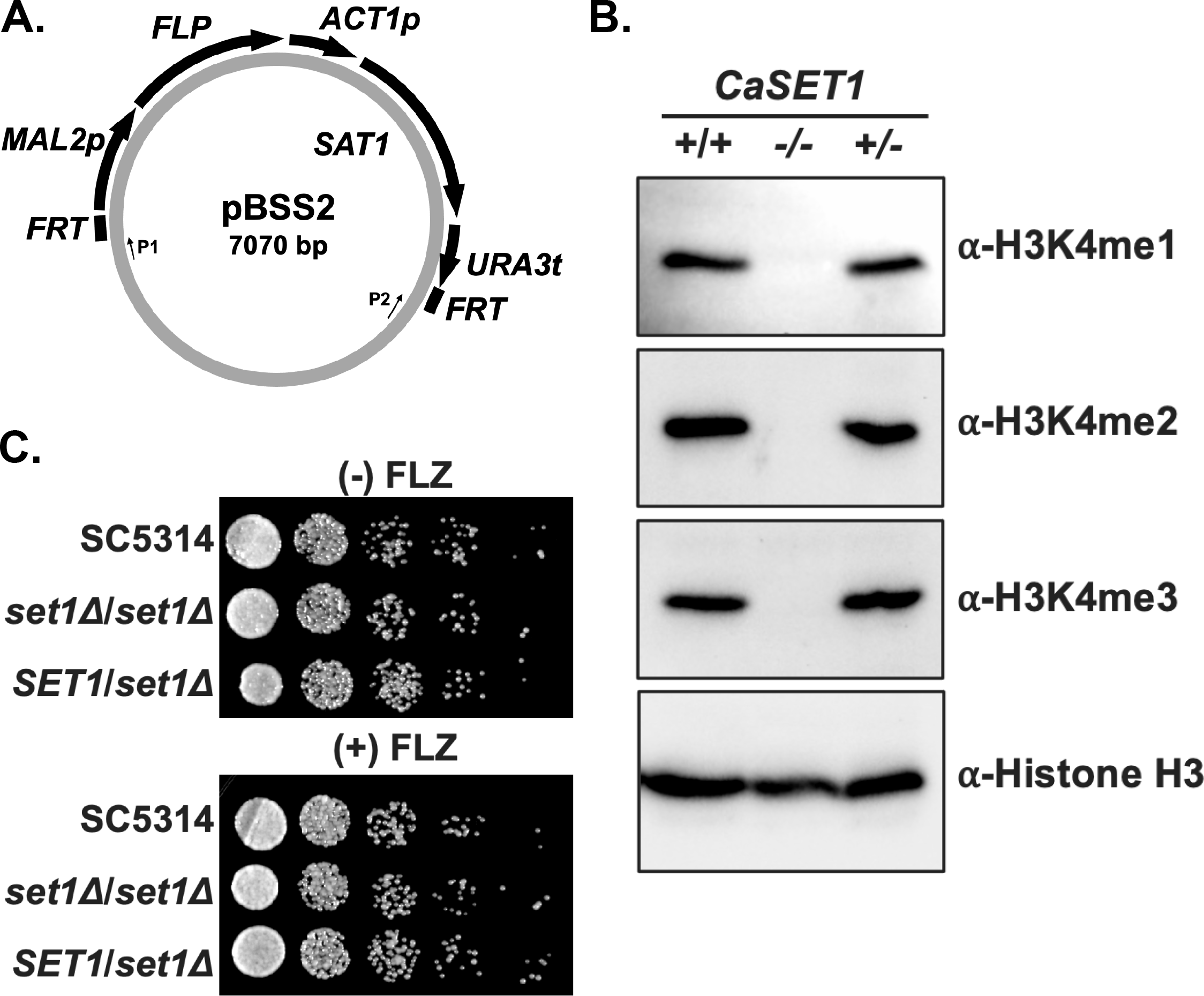
The CRISPR-Cas9 RNP system is used for deleting *SET1* in *C. albicans*. **(A)** Schematic of pBSS2-*SAT1-*FLP plasmid. P1 and P2 indicate location of amplification primer sequences. **(B)** Whole cell extracts were isolated from indicated *C. albicans* strain SC5314 and immunoblotted against methyl-specific H3K4 mono-, di- and trimethylation antibodies. Histone H3 was used as a loading control. **(C)** Five-fold serial dilution spot assays of indicated *C. albicans* strains with and without 0.5 µg/mL fluconazole (FLZ). Images were captured at 24 hours.

## DISCUSSION

In this study, we have expanded the toolkit in *Candida* by utilizing the CRIPSR-Cas9 RNP approach which allowed the repurposing of drug resistance cassettes for genetic manipulation in prototrophic strains. Using these tools, we established that deleting *ERG3* in *C. glabrata* and *C. auris* confers a fluconazole resistant phenotype. We also identified a synthetic genetic interaction between *C. glabrata ERG3* and *ERG5* and determined azole susceptible differences between *C. albicans set1Δ*/*Δ* strains and *C. glabrata* set1*Δ* strains.

We show that the use of CRISPR-Cas9 RNP with homology regions of 130-150 bp is efficient at making single gene deletions, double deletions and epitope tags in *C. glabrata*. Although three CRISPR-Cas9 RNP studies have used long flanking sequence for gene deletions in *C. glabrata* and *C. auris* (14, 15, 18), studies have shown short homology regions of ∼50-70bp are feasible for making gene deletions in *C. auris* and *C. albicans* (16, 17, 46). We have attempted to use short homology regions of ∼60 bp for deleting genes in *C. glabrata*, but it appears not to be as consistent using ∼130-150 bp flanking sequences. In addition, 130-150 bp homology regions have been useful for making double deletion and epitope tagged strains in *C. glabrata*.

Using the CRISPR-Cas9 RNP method, we also determine that two additional drug resistance cassettes (*KanMX* and *BleMX*) can be used reliably in *C. glabrata* allowing for more complex genetic manipulations. With a repertoire of four drug resistance cassettes available for use in *C. glabrata*, this greatly increases the flexibility and utility for manipulating prototrophic clinical isolates where auxotrophic makers are not readily available or feasible. In addition, our study successfully demonstrates the repurposing of *KanMX*-containing plasmids traditionally utilized for making gene deletions or C-terminal epitope tags in *S. cerevisiae*, for use in *C. glabrata*. While we clearly demonstrate that the endogenous C-terminal 3xHA tagging constructs used for *S. cerevisiae* is suitable for *C. glabrata*, this approach may not work for all genes, as C-terminal tagging may disrupt the function of the protein. Thus, our approach would also allow for repurposing endogenous N-terminal tagging constructs designed for *S. cerevisiae*. For example, our lab has generated N-ICE plasmids with *KanMX* selection cassettes for N-terminal tagging essential and non-essential genes in *S. cerevisiae* (47). We would suspect these plasmids and other *KanMX*-containing plasmids could be directly used in *C. glabrata*.

Additionally, the efficiency of endogenous epitope tagging proteins using CRISPR allows for more functional and mechanistic studies, as endogenous epitope tagged proteins have been used sparingly in prototrophic strains and clinical isolates of *C. glabrata*. This is particularly important since antibodies to endogenous proteins are scarce and costly to make.

Our study also shows that *BleMX* dominant drug selection cassette can be used in deleting genes in *C. glabrata* but also *C. auris*. Although *BleMX* has been used previously in *C. glabrata* using standard electroporation, the efficiency was extremely low and has not been typically used for routine genetic manipulation (34). *BleMX* showed the lowest efficiency of the drug selection cassettes used in our study. Nonetheless, we clearly demonstrate the CRISPR- Cas9 RNP method does improve homologous recombination efficiency enough where *BleMX* can be used. Moreover, our codon optimized *BleMX* plasmid will be readily available as another effective and needed dominant selection cassette for *C. auris*. Currently, we have not determined if our codon optimized *BleMX* drug resistance cassette can be used in other CTG clade species.

CRISPR-Cas9 RNP has been used successfully for *C. glabrata*, C*. auris*, *C. lusitaniae,* and *C. albicans* (14–18). We have also successfully used the CRISPR-Cas9 RNP approach to delete *C. albicans SET1* using a recyclable *SAT1* cassette. Interestingly, the loss of *SET1* does not confer azole susceptibility in contrast to when *SET1* is deleted in *C. glabrata* or *S. cerevisiae* which is due to either altered *ERG11* gene expression or *PDR5* expression, respectively (36, 37). Because *C. albicans* is part of the CTG clade and is more evolutionarily distant to *C. glabrata* and *S. cerevisiae*, this may suggest a species-specific utilization of *SET1*.

In contrast to *C. glabrata*, we have not been able to utilize *KanMX* in *C. albicans* due this organism’s high tolerance/resistance to the aminoglycoside antibiotic, G418. However, it has been reported that adjuvants such as quinine or molybdate can suppress background growth of *C. albicans* and allow successful integration of codon optimized *CaKan* and *CaHygB* cassettes using standard chemical transformation procedures (48). We anticipate that using these constructs, adjuvants, and the CRISPR-Cas9 RNP method could reduce background growth and increase HDR for efficient use of these markers in *C. albicans*. Alternatively, simultaneous deletion of both alleles with *KanMX* or *HygB* without adjuvants may work, since a CRISPR-RNP based system has been used successfully to simultaneously delete both alleles in *C. albicans* when using *SAT1* and *HygB* (17).

Overall, our study provides the field additional ways to efficiently manipulate *Candida* pathogens. Importantly, this approach provides us further insight in the ergosterol pathway and species differences in azole susceptibility in *Candida* pathogens when *SET1* is deleted although additional studies would be needed to further address the mechanisms of these observations.

Applying this expanded toolkit to other studies in *Candida* should enhance our understanding of fungal drug resistance and pathogenesis.

## MATERIALS AND METHODS

### Yeast strains and plasmids

All strains used are described in Table S1. *C. glabrata* strains were derived from the *Cg*2001 (ATCC 2001). *C. albicans* strains were derived from SC5314 (49), a gift from William A. Fonzi, Georgetown University. *C. auris* AR0387 strain was obtained from the CDC AR Isolate Bank. Yeast cells were grown in YPD medium or synthetic complete (SC, Sunrise Science) medium as indicated. The pAG25, pAG32, and pUG6 plasmids were obtained from Euroscarf (22, 38). The pFA6a-3HA-*KanMX* and pCY3090-07 plasmids were obtained from Addgene (35, 39). The pBSS2-*SAT1* flipper plasmid was provided to us by P. David Rogers, St. Jude Children’s Research Hospital with permission from Joachim Morschauser (43). pCdOpt-BMX (*BleMX*) was synthesized by IDT where the *TEF1p*-*BleMX-TEF1t* sequence was codon optimized for CTG clade *Candida* species using the IDT codon optimization tool, custom synthesized and cloned into the pUCIDT plasmid. The pCdOpt-BMX plasmid can be obtained at Addgene (ID number 203929).

### PCR amplification for gene deletion and epitope tagging

All oligonucleotides used are denoted in Table S2. Forward primers used for gene deletions were designed with homology regions of ∼130-150 bp flanking the 5’-ORF of the target gene of interest followed by 20-25 base pairs of sequence homologous to the indicated plasmid.

Reverse primers were designed with homology regions ∼130-150 bp flanking the 3’-ORF of the target gene of interest followed by 20-25 base pairs of sequence homology to the indicated plasmid. PCR conditions for amplification of replacement cassettes are as follows: 95°C for 5 minutes; 95°C for 30 seconds, 52°C for 30 seconds, 72°C for 2-3 minutes for a total of 30 cycles, with a final elongation step at 72°C for 10 minutes. The final PCR products were pooled and purified from agarose gels.

### CRISPR gRNA design and selection

Custom Alt-R CRISPR gRNAs were designed and ordered from Integrated DNA Technologies (Table S3). For each gene deletions, two CRISPR gRNAs were designed in close to the 5’ and 3’ ORF of the gene of interest. For epitope tagging, one CRISPR gRNA was designed in the 3’UTR of the gene of interest. CRISPR gRNAs were selected based upon their designated “On-Target Score” as determined by the CRISPR-Cas9 guide RNA design checker (IDT). Potential gRNAs were screened for Off-Target events using the CRISPR RGEN Tools Cas OFFinder (http://www.rgenome.net/cas-offinder/). Selected gRNAs required >75 On-Target Score as well as 0 potential off target events with 3 mismatches or less.

### CRISPR-Cas9 RNP system

The CRISPR-Cas9 RNP method was based on Grahl et al. with slight modifications (18). Briefly, Alt-R CRISPR crRNA and tracrRNA were used at a working concentration of 20 µM. CRISPR- Cas9 crRNAs:tracrRNA hybrid was made by mixing together 1.6 µL of crRNA (8 µM final concentration), 1.6 µL of tracrRNA (8 µM final concentration), and 0.8 µL of RNAse free water. Two crRNAs, 0.8 µL of each was added at a stoichiometric equivalent to tracrRNA and for C-terminal tagging one crRNA, 1.6 µL was used. The CRISPR RNP mix was incubated at 95°C for 5 minutes and allowed to cool to room temperature. 3 µL of 4 µM Cas9 (IDT) was added to the mix (final concentration of 1.7 µM) and incubated at room temperature for 5 minutes.

### Cell transformation

25 mL of the desired strain was grown to an OD600 of 1.6 to saturation prior to transformation. Cells were collected by centrifugation, resuspended in 10 mL 1x LiTE Buffer (100 mM LiAc, 10 mM Tris-HCl, 1 mM EDTA), and shaken at 250 rpm at 30°C for an hour. DTT was added to a final concentration of 100 mM and cells were incubated at 30°C for an additional 30 minutes. Cells were then collected by centrifugation, washed twice with 1 mL ice cold water, and washed once more with 1 mL of cold sorbitol. Cells were resuspended in 200 µL of cold sorbitol for electroporation.

### Electroporation and colony PCR

20 µL of prepared cells, 1-3 µg of drug resistant cassette DNA, CRISPR mix, and RNAse free water to a final volume of 45 µL was mixed and transferred to a BioRad Gene Pulser cuvette (0.2 cm gap). Cells were pulsed using an Eppendorf Eporator at 1500 V and immediately resuspended in 1 mL of ice-cold Sorbitol. Cells were then collected by centrifugation, resuspended in 1 mL of YPD media, and allowed to recover by incubation at 30°C at 250 rpm for 3-24 hours. Cells were then collected, resuspended in 100 µL of YPD, and plated onto drug selective media at the desired concentration. Nourseothricin (GoldBio) was used at a final concentration of 300 µg/mL for antibiotic selection of the *NatMX* cassette. Hygromycin B (Cayman) was used at a final concentration of 500 µg/mL antibiotic selection of the *HphMX* cassette. Geneticin (G418, GoldBio) was used at a final concentration of 800 µg/mL for antibiotic selection of the *KanMX* cassette. Zeocin (Cayman) was used at a final concentration of 600 µg/mL for antibiotic selection of the *BleMX* cassette in *C. glabrata* and 800 µg/mL in *C. auris*. Colonies were streaked onto fresh plates with the desired drug, and single colonies were selected and restreaked onto fresh YPD plates. Colonies were screened via colony PCR using primers indicated in Table S2. Three independent clones were verified by PCR and analyzed for phenotypic characterizations.

### Serial dilution growth assay

For serial dilution spot assays, yeast strains were inoculated in SC media and grown to saturation overnight. Yeast strains were diluted to an OD600 of 0.1 and grown in SC media to log phase shaking at 30°C. The indicated strains were spotted in five-fold dilutions starting at an OD600 of 0.01 on untreated SC plates or plates containing 8 µg/mL, 16 µg/mL, or 64 µg/mL fluconazole (Cayman). For *C. glabrata* and *C. auris*, plates were grown at 30°C on SC plates for 48 hours prior to imaging. For *C. albicans*, plates were grown at 30°C on YPD plates for 48 hours prior to imaging.

### Quantitative real-time PCR analysis

RNA was isolated from cells grown in SC media by standard acid phenol purification as previously described (37). Reverse transcription to generate cDNA was performed using the ABM All-In-One 5X RT MasterMix (ABM). Primer Express 3.0 software was used for designing primers for gene expression analysis by quantitative real-time polymerase chain reaction (Table S4). A minimum of three biological replicates, as well as three technical replicates, were performed for each biological replicate. All data were analyzed using the comparative CT method (2^-ΔΔCT^). *RDN18* (18S ribosomal RNA) was used as an internal control. All samples were normalized to the *Cg*2001 WT strain (Table S5).

### Cell extract and Western blot analysis

Whole cell extraction and Western blot analysis were performed as previously described (50, 51). The anti-HA (Roche 12CA5, 1:10,000) monoclonal antibody was used as previously described (52). Histone H3K4 methylation-specific antibodies were used as previously described: H3K4me1 (Upstate, 07-436; 1:2,500), H3K4me2 (Upstate, 07-030; 1:10,000), and

H3K4me3 (Active Motif 39159, 1:100,000) (42, 53). Histone H3 rabbit polyclonal antibody (PRF&L) was used at a 1:100,000 dilution.

## ACKNOWLEGEMENTS

This publication was supported by grants from the National Institute of Allergy and Infectious Diseases of the National Institutes of Health under award number T32AI148103 (To J.B.G.) and AI136995 (To S.D.B.). Funding support was also provided by the NIFA 1007570 (To S.D.B) and NSF DBI-2150331 (To M.G.B). We thank Drs. Majid Kazemian and Mark Hall for critical review of our manuscript.

## Supplemental Material

### SUPPLEMENTAL TABLES

**Table S1:** Yeast Strains and Genotype

| Yeast Strain | Genotype | Reference | Strain Name |
| --- | --- | --- | --- |
| ATCC 2001<br>*Candida glabrata* WT | *C. glabrata* prototrophic wild type strain | www.atcc.org | CgWT |
| SDBY1620 | *Cg2001 erg3Δ::NatMX* | This study | *Cgerg3Δ NatMX* |
| SDBY1621 | *Cg2001 erg3Δ::HphMX* | This study | *Cgerg3Δ HphMX* |
| SDBY1622 | *Cg2001 erg3Δ::KanMX* | This study | *Cgerg3Δ KanMX* |
| SDBY1623 | *Cg2001 erg3Δ::BleMX* | This study | *Cgerg3Δ BleMX* |
| SDBY1624 | *Cg2001 erg5Δ::NatMX* | This study | *Cgerg5Δ* |
| SDBY1625 | *Cg2001 erg3Δ::NatMX erg5Δ::HphMX* | This study | *Cgerg3Δerg5Δ* |
| SDBY1626 | *Cg2001 ERG3-3xHA::KanMX* | This study | *CgERG3-3xHA* |
| SDBY1627 | *Cg2001 ERG11-3xHA::KanMX* | This study | *CgERG11-3xHA* |
| SC5314 | *C. albicans* prototrophic wild type strain | www.atcc.org | SC5314 |
| SDBY1628 | SC5314 *SET1/set1Δ::FRT* | This study | *CaSET1/set1Δ* |
| SDBY1629 | SC5314 *set1Δ::FRT/set1Δ::FRT* | This study | *Caset1Δ/set1Δ* |
| *C. auris* AR0387 | *C. auris* clinical isolate | CDC AR Isolate Bank | *Caur0387* |
| SDBY1630 | *Caur0387 erg3Δ::BleMX* | This study | *Caur0387erg3Δ* |

**Table S2:**
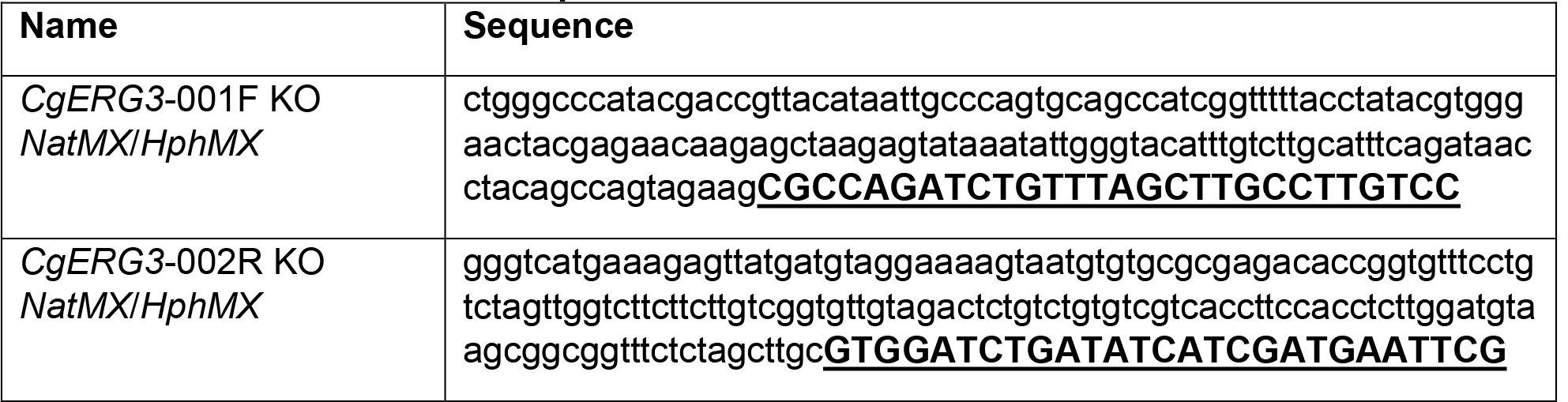
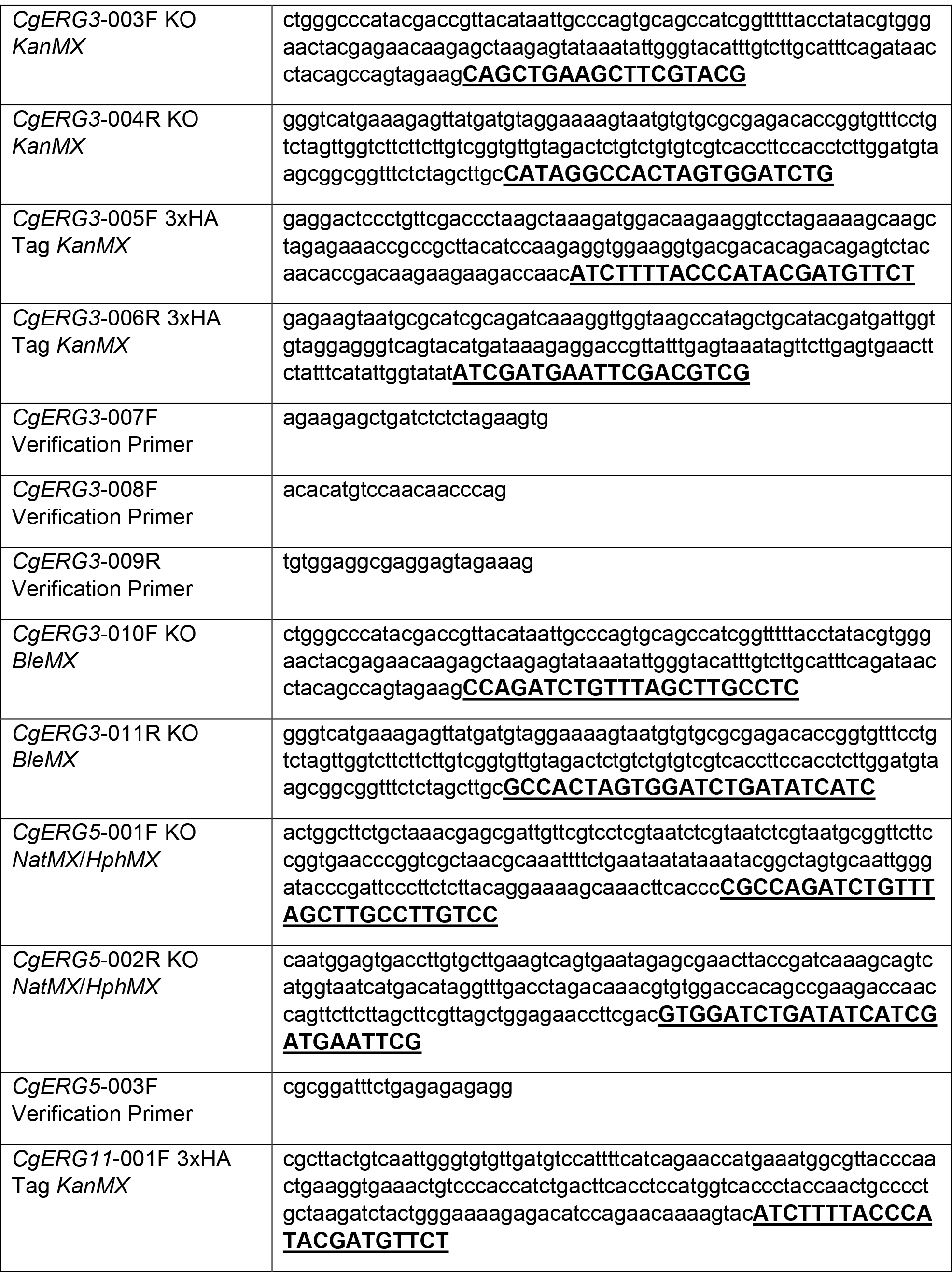
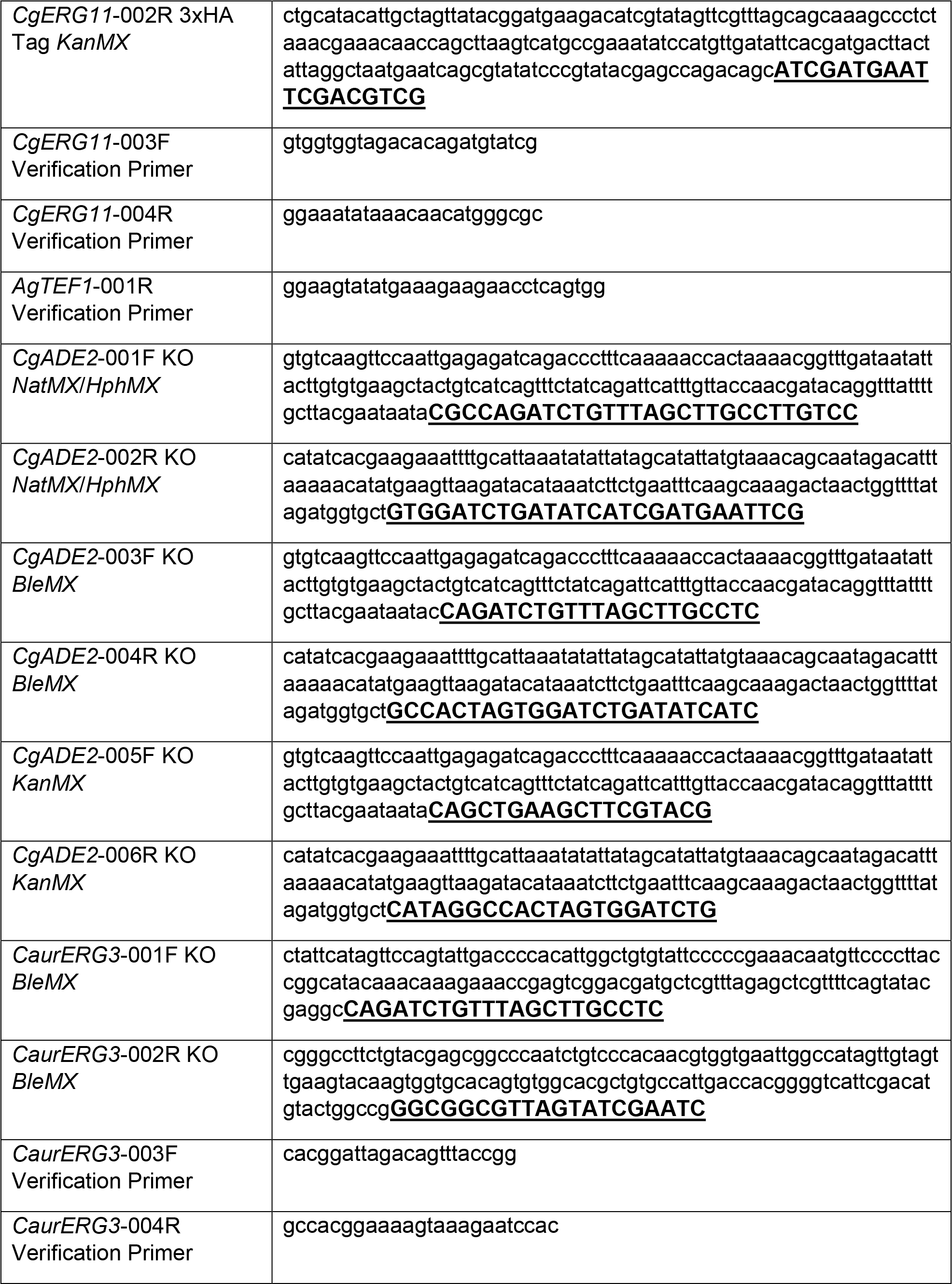
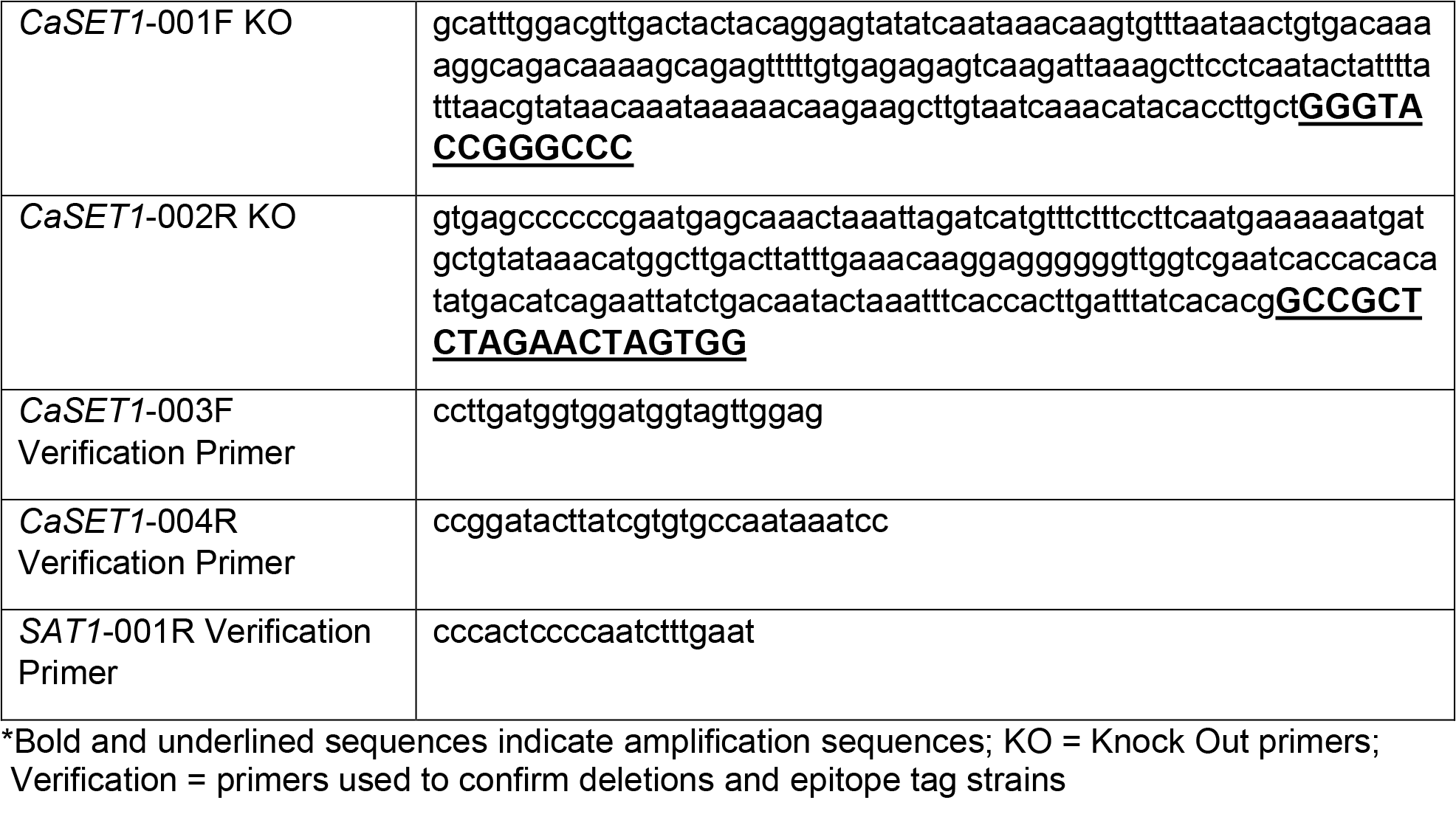
Primer Names and Sequences

**Table S3:**
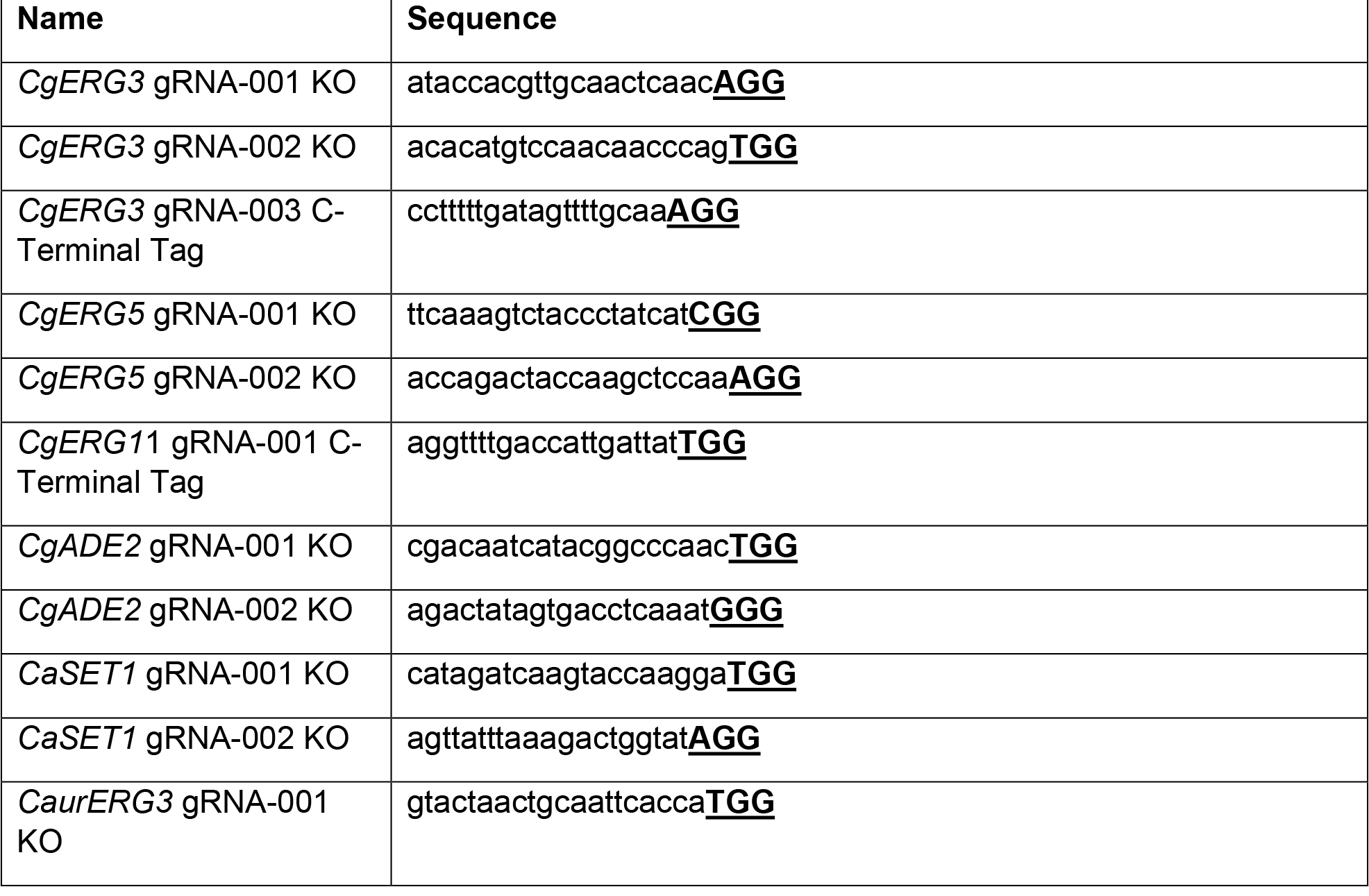
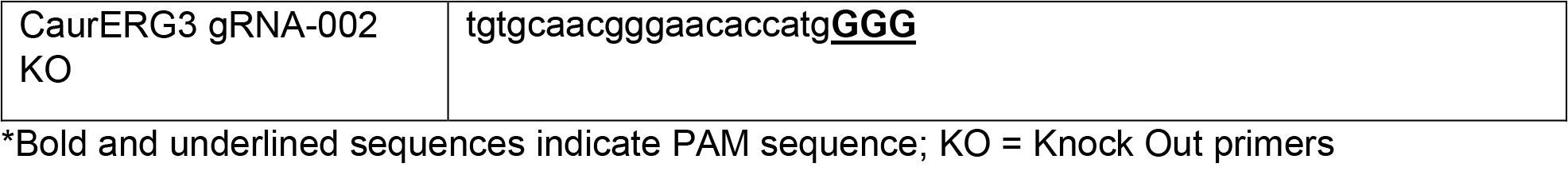
CRISPR gRNA sequences

**Table S4:**
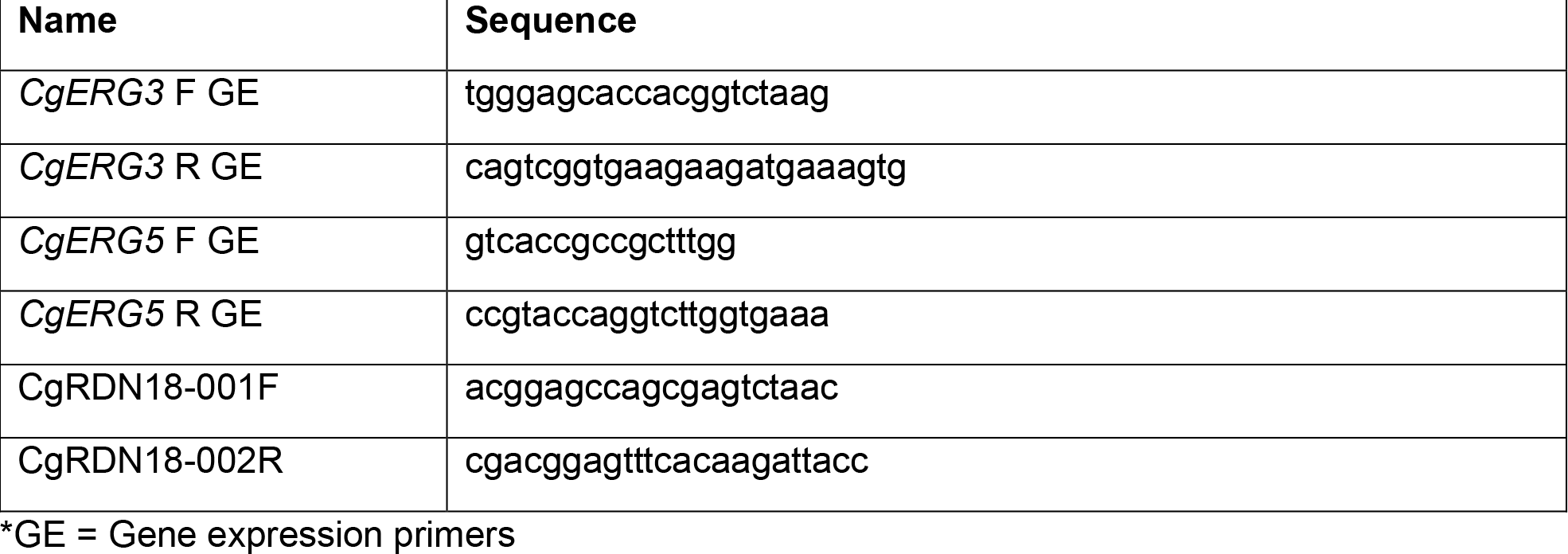
qRT-PCR Primers

**Table S5:** qRT-PCR

| Gene | Strain | Mean RQ | St. Dev | N |
| --- | --- | --- | --- | --- |
| *CgERG3* | *CgWT* | 1.00 | 0.42 | 3 |
| *CgERG3* | *Cgerg3Δ* | 0.00 | 0.00 | 3 |
| *CgERG3* | *Cgerg3Δerg5Δ* | 0.00 | 0.00 | 3 |
| *CgERG3* | *Cgerg5Δ* | 0.38 | 0.22 | 3 |
| *CgERG5* | *CgWT* | 1.00 | 0.61 | 3 |
| *CgERG5* | *Cgerg3Δ* | 2.17 | 0.79 | 3 |
| *CgERG5* | *Cgerg3Δerg5Δ* | 0.00 | 0.00 | 3 |
| *CgERG5* | *Cgerg5Δ* | 0.00 | 0.00 | 3 |

